# New guidelines for DNA methylome studies regarding 5-hydroxymethylcytosine for understanding transcriptional regulation

**DOI:** 10.1101/334318

**Authors:** Le Li, Yuwei Gao, Qiong Wu, Alfred S. L. Cheng, Kevin Y. Yip

## Abstract

Many DNA methylome profiling methods cannot distinguish between 5-methylcytosine (5mC) and 5-hydroxymethylcytosine (5hmC). Since 5mC typically acts as a repressive mark whereas 5hmC is an intermediate form during active demethylation, the inability to separate their signals could lead to incorrect interpretation of the data. Meanwhile, many analysis pipelines quantify methylation level by the count or ratio of methylated reads, but the proportion of discordant reads (PDR) has recently been proposed to be a better indicator of gene expression level. Is the amount of extra information contained in 5hmC signals and PDR worth the additional experimental and computational costs? Here we combine whole-genome bisulfite sequencing (WGBS) and oxidative WGBS (oxWGBS) data in normal human lung and liver tissues and their paired tumors to investigate the quantitative relationships between gene expression and signals of the two forms of DNA methylation at promoters, transcript bodies, and immediate downstream regions. We find that 5mC and 5hmC signals correlate with gene expression in the same direction in most samples, but considering both types of signals increases the accuracy of expression levels inferred from methylation data by a median of 18.2% as compared to having only standard WGBS data, showing that the two forms of methylation provide complementary information about gene expression. In addition, differential analysis between matched tumor and normal pairs is particularly affected by the superposition of 5mC and 5hmC signals in WGBS data, with at least 25-40% of the differentially methylated regions (DMRs) identified from 5mC signals not detected from WGBS data. We do not find PDR to be more informative about expression levels than ratio of methylated reads, and integrating the two types of methylation features only improves the accuracy of inferred expression levels by at most 9.8%. Our results also confirm previous finding that methylation signals at transcript bodies are more indicative of gene expression levels than promoter methylation signals, and further show that in addition to the first exon, methylation signals at the last exon and internal introns also contain non-redundant information about gene expression. Overall, our study provides concrete data for evaluating the cost effectiveness of some experimental and analysis options in the study of DNA methylation in normal and cancer samples.

## Introduction

DNA methylation, the methylation of the carbon 5 atom of cytosines, usually occurs within the CpG context in eukaryotes, and in some cell types also CpHpG and CpHpH contexts (Bird, 2002; Cokus et al., 2008; Lister et al., 2009). It is involved in various biological processes, including embryonic development, genomic imprinting, X chromosome inactivation and genome stability maintenance (Lister et al., 2009). Aberrant DNA methylation is associated with a variety of human diseases, including cancer (Robertson, 2005). Many types of cancer exhibit global hypomethylation as compared to normal tissues, while specific loci could be hypermethylated (Ehrlich, 2009; Klutstein et al., 2016).

DNA methylation is tightly related to gene expression. Methylation of CpG islands within promoter regions is associated with long-term gene silencing, while methylation in other regions are more dynamic and tissue-specific (Jones, 2012). Sequences up to 2kb away from CpG islands, termed CpG island shores, have been shown to display differential methylation in cancer that correlates with differential gene expression (Irizarry et al., 2009). DNA methylation at regulatory elements other than promoters is less studied, but some recent work has started to demonstrate correlations between enhancer methylation and gene silencing (Aran et al., 2013; Cao et al., 2017; Heyn et al., 2016). How gene body methylation is related to gene expression has been more controversial, with a positive correlation between them observed in some cell types but not in some others. Mechanistically, gene body methylation could be related to repression of anti-sense transcript, efficiency of transcription elongation, usage of alternative promoter, and RNA splicing (Choi et al., 2009; Lorincz et al., 2004; Maunakea et al., 2010; Rountree and Selker, 1997).

The ambiguous relationship between gene body methylation and gene expression could be partly due to the presence of multiple forms of DNA methylation. 5-methylcytosine (5mC) can be converted by the Ten-eleven translocation (Tet) family of proteins into 5-hydroxymethylcytosine (5hmC), 5-formylcytosine (5fC), and 5-carboxylcytosine (5caC) during active demethylation (Song et al., 2012). Unlike 5mC’s characteristic enrichment at promoters of repressed genes, 5hmC has been found enriched at active enhancers and around expressed genes, including gene body regions (Song et al., 2012; Yu et al., 2012). If an experimental method cannot distinguish between 5mC and 5hmC (or the other two intermediate forms), depending on their relative levels, methylation may appear to correlate with gene expression in different ways for different genes with the same total methylation level.

Unfortunately, that is exactly the situation with many commonly used experimental methods. In standard bisulfite conversion, which is used in whole-genome bisulfite sequencing (WGBS), reduced representation bisulfite sequencing (RRBS) and Infinium 27k/450k/EPIC arrays, both 5mC and 5hmC are unconverted. The resulting data can only tell whether a cytosine is methylated or not, but cannot tell which form of methylation a cytosine takes (Jin et al., 2010). Some other methods such as MeDIP-seq and MBDCap-seq could specifically detect 5mC (Jin et al., 2010), but they do not offer single-base resolution and fail to provide information about the other forms of DNA methylation.

Recently, realizing the importance of 5hmC as a representative of the demethylation forms, a number of methods have been proposed to detect it at single-base resolution (Li et al., 2016; Petterson et al., 2014). Some of these methods are based on oxidative bisulfite sequencing (oxBS), which specifically detects 5mC. Conceptually, by subtracting the methylation level detected by oxBS from that detected by standard bisulfite sequencing (BS), the 5hmC level can also be deduced. In practice, this subtraction could lead to negative values due to errors and stochastic factors in the experimental and analysis procedures, which can be corrected computationally (Xu et al., 2016).

Currently, it is still not completely clear how 5mC and 5hmC at promoter, gene body and downstream regions are related to gene expression, both separately and jointly. For instance, was the previously observed positive correlation between gene body methylation and gene expression purely due to 5hmC? Do 5mC and 5hmC correlate with gene expression in opposite directions? If 5mC level is already known, does 5hmC level provide extra information about gene expression, in normal and disease samples?

Besides, a lot of existing knowledge about DNA methylation is qualitative rather than quantitative. For example, strong promoter 5mC is known to be associated with gene silencing, but the expected amount of gene expression given a certain level of promoter methylation is usually not known. More generally, if the 5mC and 5hmC levels at the promoter, gene body and downstream region are measured, is it possible to tell the corresponding expression level of a gene? This question is particularly important in epigenenomic studies of diseases, in which a common question is whether an observed expression level change of a gene can be attributed to the promoter methylation change alone or it is also affected by methylation at other regions or even by other regulatory mechanisms.

In addition to the methylation level (“beta value”), defined as the proportion of methylated reads/signal intensity at a CpG site among the total number of reads/signal intensity, it has been proposed that the proportion of discordant reads (PDR), defined as the ratio of reads having discordant methylation status at different CpGs, is a better indicator of gene expression level in chronic lymphocytic leukemia (Landau et al., 2014). Is PDR generally more informative than beta values, especially when signals for 5mC and 5hmC are separately measured?

In this work, we use WGBS and oxWGBS data from normal liver and lung tissues and paired cancer samples to study the quantitative relationships between gene expression and the two DNA methylation forms, 5mC and 5hmC, quantified by both beta values and PDR, at different genic and regulatory elements associated with the transcripts. In addition to answering the above conceptual questions, another goal of this study is to provide practical guidelines as to whether both 5mC and 5hmC should be measured and whether both beta values and PDR should be computed, neither of which is a common practice currently.

## Results

### 5mC and 5hmC levels alone can partially infer transcript expression level

We obtained WGBS, oxWGBS and RNA sequencing (RNA-seq) data for 12 samples, including three pairs of human normal liver tissues (Liver N1-N3) and matched tumors (Liver T1-T3), and three pairs of human normal lung tissues (Lung N1-N3) and matched tumors (Lung T1-T3) (Li et al., 2016). Based on these data, for each transcript, we computed the average raw WGBS and oxWGBS beta values, as well as the inferred 5mC and 5hmC levels at its 16 associated upstream, transcript body and downstream regions in each sample (Figure 1a, Materials and Methods). Heat maps of the resulting data set reveal some subtle correlations between these methylation features at the different associated regions and the corresponding expression levels of the transcripts, with lower methylation at promoters and some body features for transcripts with higher expression (Figures 1b, S1). A hierarchical clustering of the samples based on all their methylation features shows two main clusters corresponding to the two tissues of origin rather than cancer status (Figure 1c). A related observation has recently been made based on gene expression data of 8,000 patients of 17 cancer types, that liver tumors are more similar to normal liver tissues than to other types of tumors (Uhlen et al., 2017).

**Figure 1:**
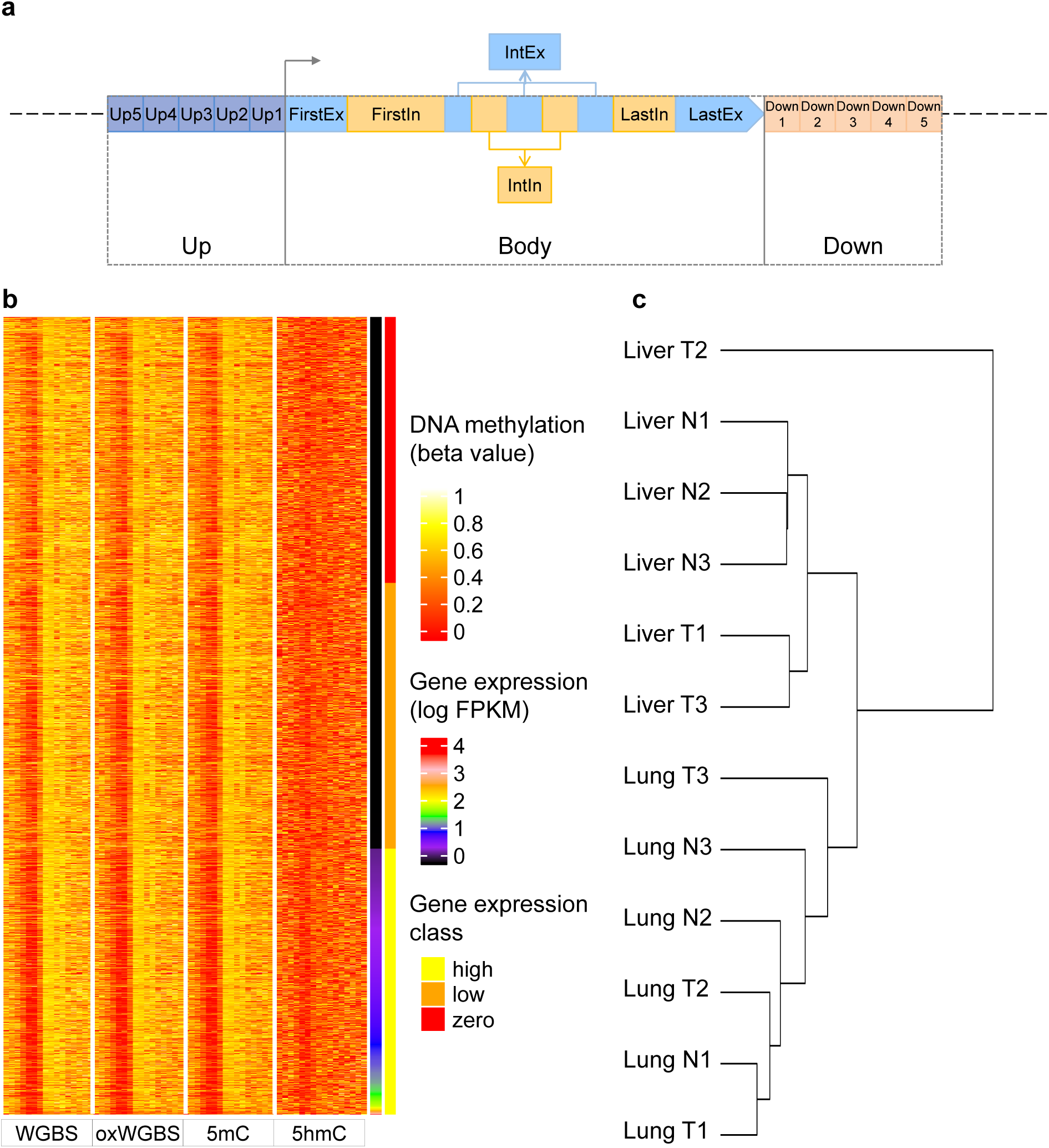
Definition of the regions associated with each transcript and the resulting data set. a Genomic regions defined for each transcript at which the beta values or PDR values were used to infer expression level of the transcript. Both the upstream (Up) and downstream (Down) regions were divided into five 400bp bins (Up1-Up5 and Down1-Down5). The transcript body (Body) was divided into first exon (FirstEx), first intron (FirstIn), internal exons (IntEx), internal introns (IntIn), last exon (LastEx) and last intron (LastIn). b A heat map of the resulting large data set for sample Liver T1. Each row represents a transcript and the transcripts are sorted in ascending order according to their expression levels. The four blocks of columns represent beta values based on WGBS, oxWGBS, 5mC and 5hmC, respectively. Within each block, the different columns are respectively Up5-Up1, FirstEx, FirstIn, IntEx, IntIn, LastEx, LastIn and Down1-Down5. After the four methylation blocks, the last two columns show the log expression level and expression class, respectively. c Hierarchical clustering of the samples based on all their methylation features in the large data set using Ward’s method. 21

We also defined a similar data set with both beta values and PDR values computed. While beta values can be computed from the raw WGBS and oxWGBS data as well as the processed 5mC and 5hmC levels, PDR values can only be defined directly from sequencing reads, and were thus computed from raw WGBS and oxWGBS data only. To ensure reliable calculations of PDR values, only regions with sufficient read coverage were considered (Materials and Methods), leading to a smaller number of transcripts included in this data set. Hereafter, we refer to this data set as the “small” set and the data set with only beta value features as the “large” set. In the followings we first focus on the analyses of the large data set.

To investigate whether beta values in the regions associated with a transcript are indicative of its expression level, we performed statistical modeling of transcript expression classes (zero-, low-and high-expression), and evaluated the accuracy of the resulting models using a rigorous cross-validation procedure (Materials and Methods).

Considering methylation levels in all 16 regions associated with each transcript, the constructed models were fairly accurate in separating transcripts belonging to the different expression classes, with a median AUROC (area under the receiver-operator characteristic) of around 0.7 (Figure 2a, “BS+oxBS+5mC+5hmC”). This value is close to the AUROC reported in a previous study that involved only WGBS features (Lou et al., 2014) despite a more rigorous evaluation procedure and a different way of quantifying methylation level used in the current study. Comparing the different expression classes, the DNA methylation features were more successful in identifying transcripts with zero or high expression than those with an intermediate expression level (Figure S2).

**Figure 2:**
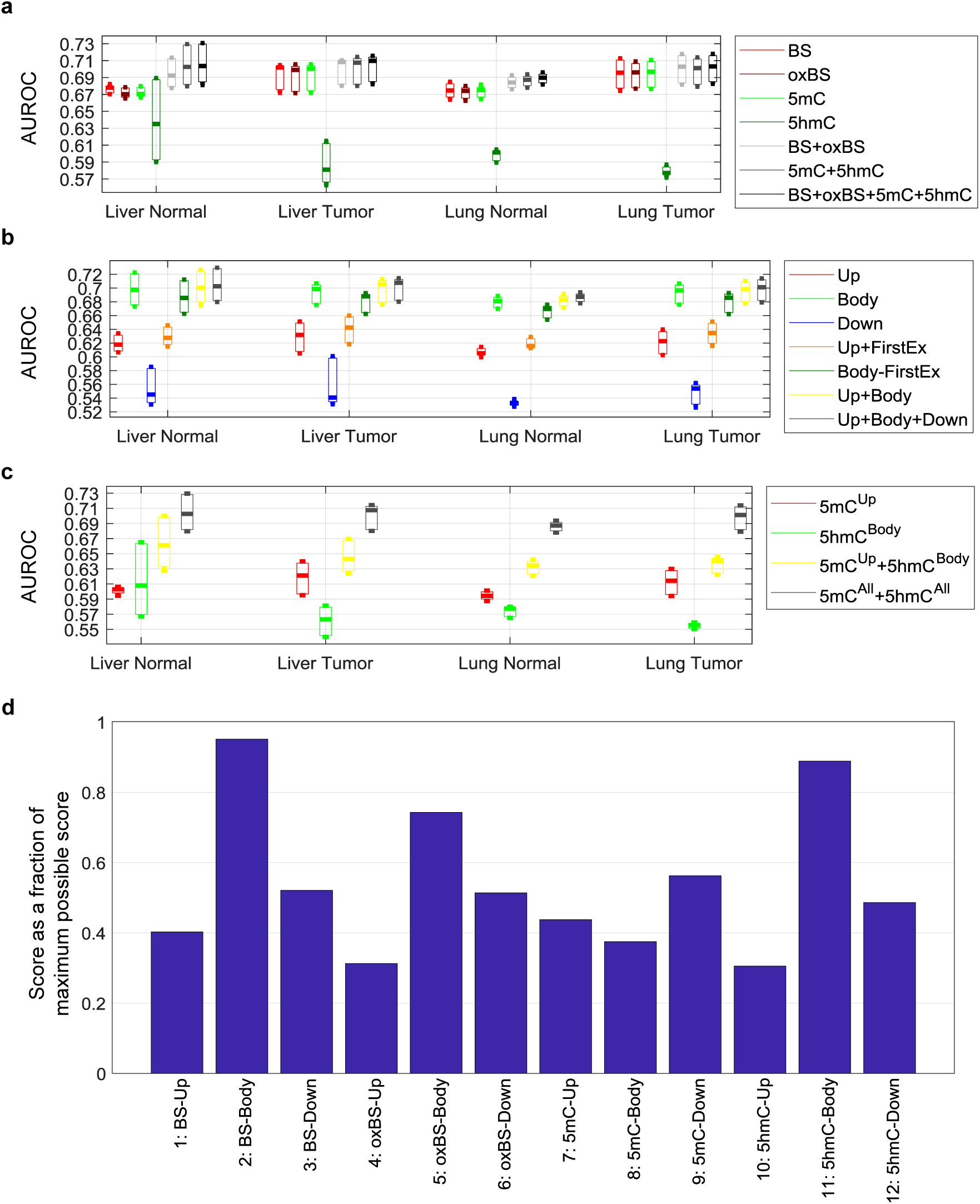
Accuracy of the models for inferring expression classes based on the large data set. a-c Each bar represents the distribution of AUROC values across the three expression classes of the three samples in each sample group. a Comparison of models involving different combinations of methylation features from all associated genomic regions of the transcripts. b Comparison of models involving both 5mC and 5hmC levels at different combinations of genomic regions associated with each transcript. c Comparison of several knowledge-driven models. d The most useful methylation feature blocks for inferring gene expression level based on the forward-search procedure of feature selection. For each sample, the top feature block was given a score of 8, the second given a score of 7, and so on, for the top 8 feature blocks. The total score of each feature block across all 12 samples is shown as a percentage of the maximum possible score of 8×12=96.

To evaluate whether these results are sensitive to the transcript annotation set, we repeated the above procedures considering only protein-coding transcripts and/or only the transcripts with experimental evidence or manual curation (from Gencode levels 1 and 2). The resulting AUROC values were similar for all these settings (Figure S3), suggesting that the models constructed were general for both protein-coding and non-coding genes, and for transcripts at different confidence levels.

### 5mC and 5hmC provide complementary information about gene expression

We then investigated the relative importance of the DNA methylation features by constructing models using only subsets of features. First, we compared methylation features derived from the four types of methylation data considering all 16 associated regions (Figure 2a). Among the models involving only features derived from a single type of data, the models with WGBS, oxWGBS or 5mC features had similar AUROC values, all higher than models with 5hmC features. On the other hand, models involving both 5mC and 5hmC features (“5mC+5hmC”) performed better than models involving either 5mC or 5hmC features alone, most substantially for the normal liver samples, showing that these two forms of DNA methylation provide complementary information about gene expression. Similarly, combining both WGBS and oxWGBS features (“BS+oxBS”) slightly improved the modeling results as compared to having only WGBS or only oxWGBS features. Finally, models involving all four types of data (“BS+oxBS+5mC+5hmC”) had similar performance as models involving only the inferred 5mC and 5hmC levels (“5mC+5hmC”), suggesting that these derived DNA methylation features successfully captured the essential information about gene expression contained in the raw WGBS and oxWGBS data. Overall, models involving all features had a median AUROC improvement of 0.7-7.2% for the different samples as compared to the models involving only WGBS features.

Next, we compared models involving methylation features at the different regions associated with each transcript (Figure 2b). Among the upstream, transcript body and downstream features, features at the transcript body were most indicative of the expression class, followed by those at the upstream regions. The higher accuracy of the transcript body models was partially, but not completely, due to the effect of the first exon, in that including the first exon always led to better modeling accuracy (comparing “Up+FirstEx” with “Up”, and comparing “Body” with “Body-FirstEx”), but models involving transcript body features were consistently more accurate than those involving upstream features no matter first exon was included or excluded in both sets (comparing “Body” with “Up+FirstEx”, and comparing “Body-FirstEx” with “Up”). Integrating features in both transcript body and upstream regions (“Up+Body”) or all three region types (“Up+Body+Down”) only improved the modeling accuracy slightly as compared to the models involving transcript body features alone.

It is well-accepted that high 5mC level at promoters is an indicator of gene repression (Miranda and Jones, 2007; Suzuki and Bird, 2008), while 5hmC has been shown to be associated with gene bodies (Stroud et al., 2011). We checked whether these knowledge-driven features are redundant and whether together they are sufficient for inferring gene expression level to the maximal accuracy. We found that combining 5mC features at upstream regions and 5hmC features at transcript bodies indeed improved modeling accuracy as compared to having either set of features alone, but their combination was still not sufficient to reach the accuracy of models involving all types of methylation features at all associated regions of the transcripts (Figure 2c), suggesting that methylation features other than promoter 5mC and transcript body 5hmC levels also contribute substantially to the understanding of transcript expression levels.

To make sure that the above observations are not specific to our definition of expression classes, we also constructed regression models to infer log expression levels of transcripts directly. The resulting correlation values (Figure S4) displayed trends highly similar to the AUROC values from the classification models, thereby confirming the generality of the results. For example, combining 5mC and 5hmC features led to better results than having either alone (Figure S4a,b) and transcript body features could infer expression levels more accurately than upstream features (Figure S4c,d).

Comparing models involving only WGBS features with those involving all methylation features (Figure S4a, “BS” vs. “BS+oxBS+5mC+5hmC”), the median Pearson correlation (across the 12 samples) between the predicted and actual log expression values increased from 0.21 to 0.25, which is equivalent to a 18.2% improvement. Among the two tissue types, liver samples had a larger increment of 26.4%.

### Feature importance and the smallest set of features with maximal information about gene expression

To systematically determine the most important methylation features for explaining expression variability, we defined a feature importance score based on the frequency of each feature being selected as one of the top features in a forward-searching procedure (Materials and Methods). When we grouped features into upstream, transcript body and downstream feature blocks (Figure 2d), WGBS and 5hmC signals at transcript bodies received the highest importance scores. This is particularly interesting because although 5hmC features alone could not infer expression levels accurately, they provided the best complementation to the WGBS features while other individually strong features (such as 5mC-Body) appeared to provide less extra information not already contained in WGBS features at transcript bodies. Among the upstream features, as expected 5mC was selected as most important.

We then further studied the 16 individual regions (Figure S5), and found that in addition to first exon (“FirstEx”) and the upstream region closest to the TSS (“Up1”), which are important for transcription factor binding and transcription initiation, some other features also consistently showed up among the most important features, including the last exon (“LastEx”) and internal introns (“IntIn”). These regions may affect transcription through other independent mechanisms such as transcriptional elongation and splicing, and were therefore selected as the next most important features.

Using the models involving all features as the best case, we investigated how the model accuracy changed as we included each additional feature or feature block during the forward searching procedure. In terms of feature blocks (Figure S6), usually 3-4 blocks were sufficient to reach the best-case performance, and these top blocks were predominantly transcript body features. In terms of individual methylation features (Figure S7), usually 10 or more features were necessary to reach the best-case performance. Although the first 3-4 top features, mainly form transcript bodies, provided the most rapid improvement of modeling accuracy, the remaining 6-8 features still provided non-negligible improvements, and sometimes they also included upstream and downstream features.

### The constructed models remain reasonably accurate when applied to other samples

All the results described above were obtained by training and testing on distinct subsets of transcripts from the same sample using a cross-validation procedure. This procedure was designed to avoid overfitting the training data, such that the models could capture the general relationships between DNA methylation and gene expression rather than trends specific to the training sample only. To confirm this generality, we applied models trained on a subset of transcripts from a sample to infer the expression class of a different subset of transcripts in another sample. The results (Figure 3) reveal that except for normal liver samples that appear to be more distinct from the other samples, our constructed models could infer expression classes of transcripts in other samples as accurately as in the training sample, which can be seen by having off-diagonal AUROC values in the result matrix not substantially lower than the diagonal ones. Importantly, a large portion of these models did not exhibit strong tissue-or disease state-specificity, with similar AUROC values no matter the testing sample had the same tissue type or disease state as the training sample or not.

**Figure 3:**
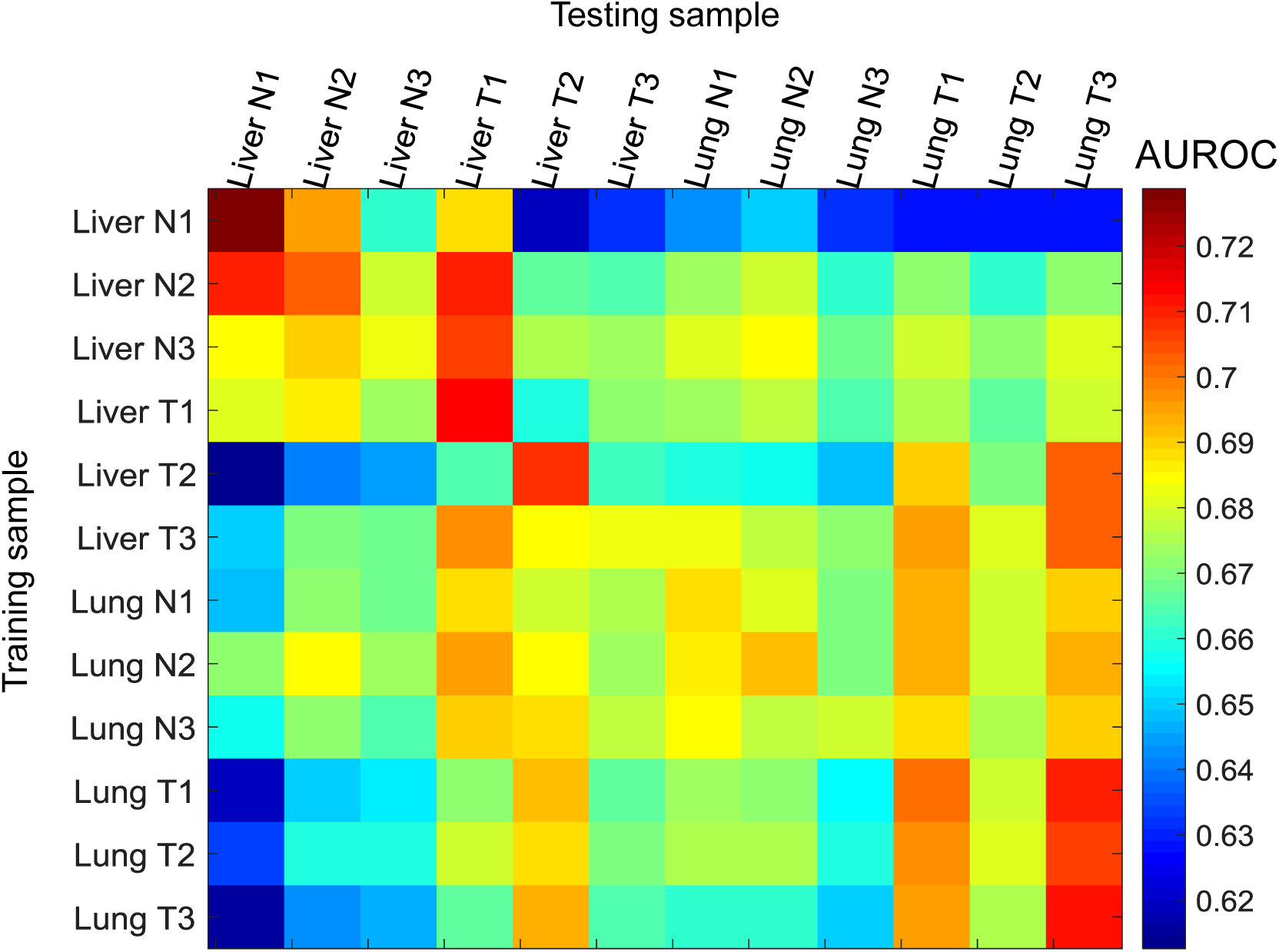
Generality of the models for predicting expression classes with all methylation features based on the large data set. Each row corresponds to a sample from which the model was trained, and each column represents a sample to which the model was applied, based on which the evaluation measure was computed. The training and testing transcripts were disjoint no matter the testing sample was the same as or different from the training sample.

### Relationship between transcript body methylation and expression

Whether DNA methylation at transcript body correlates positively or negatively with gene expression has been controversial (Ball et al., 2009; Lister et al., 2009; Lou et al., 2014; Rauch et al., 2009). Besides differences in measuring and quantifying methylation levels in the previous studies that could have led to discrepancies in their results, it has not been clear whether the relative 5mC and 5hmC levels could also be a key factor since many of these studies did not consider the two forms of DNA methylation separately. From our data, we found that at regions closest to the TSS (Up1 and FirstEx), both 5mC and 5hmC features correlated negatively with gene expression, although the correlation was stronger for 5mC (Figure S8). This is consistent with a recent report that both types of DNA methylation could repress gene expression by affecting transcription factor binding at the promoter (Kitsera et al., 2017). Inside the transcript body, 5hmC tended to correlate more positively with gene expression than 5mC in normal liver samples, but the reverse is observed for some liver and lung tumor samples. We further explored the joint relationships of 5mC and 5hmC levels at transcript body with gene expression, treating the whole transcript body as a single region (Figure S9). We did not observe any clear trends, except that highly expressed transcripts usually did not have very high 5mC or 5hmC levels at their bodies. These results reiterate that transcript body methylation has more subtle relationships with gene expression than promoter methylation.

### Comparisons between beta value and PDR features

After exploring properties of 5mC and 5hmC levels using the large data set, we then switched to the small data set to study PDR values (Figure S10). Overall, the AUROC values were generally lower than those obtained from the large data set (Figure 2), likely due to the much smaller number of transcripts in the small data set that forbade reliable modeling. Nevertheless, this data set still allowed us to explore the relative importance of beta value and PDR features. Comparing the models involving these two types of features (Figure S10a), models involving beta value features had higher AUROC values no matter WGBS, oxWGBS or both types of data were used. Combining the two types of features resulted in more accurate models in all cases. Compared with having beta value features alone, incorporating PDR features led to 1.0-5.2%, 0.6-3.7% and 0.0-3.5% AUROC improvements across the samples when WGBS, oxWGBS or both types of data were used, respectively. The same trends were also obtained from the regression results (Figure S11), with the median Pearson correlation improved up to 9.5%, 9.8% and 4.8% by incorporating PDR features when WGBS, oxWGBS or both types of data were used, respectively.

Since PDR values could only be computed from the raw WGBS and oxWGBS data, we checked whether it would be beneficial to also incorporate beta value features of the derived 5mC and 5hmC levels. The results (Figure S10b) show that adding these features (“(BS+oxBS)_Beta+PDR_+(5mC+5hmC)_Beta_”) only led to a small increase of AUROC in normal liver samples as compared to not adding them (“(BS+oxBS)_Beta+PDR_”), and did not lead to any clear improvements in other samples. These results again show that it is sufficient to define beta value features using either the raw WGBS and oxWGBS data alone or the processed 5mC and 5hmC data alone.

### Necessity of integrating 5mC and 5hmC in differential analyses

The presence of both tumor and matched normal samples in our data enabled us to investigate the necessity of measuring both 5mC and 5hmC in studying differential methylation in cancer. We first checked whether differential expression class could be inferred by beta value features (Materials and Methods). The results (Figure 4a) show that, as expected, transcripts with strong differential expression were more easily identified than those with only weak differential expression. In general, the beta value features were more successful in detecting differentially expressed transcripts in the liver sample pairs than in the lung sample pairs (Figure 4a), and when the four classes contained transcripts with more distinct differential expression profiles (Figure S12a). Again, combining both 5mC and 5hmC data led to the best modeling accuracy (Figure S12b), and methylation levels at transcript bodies were more useful than those at promoters or downstream regions in inferring the differential expression classes (Figure S12c).

**Figure 4:**
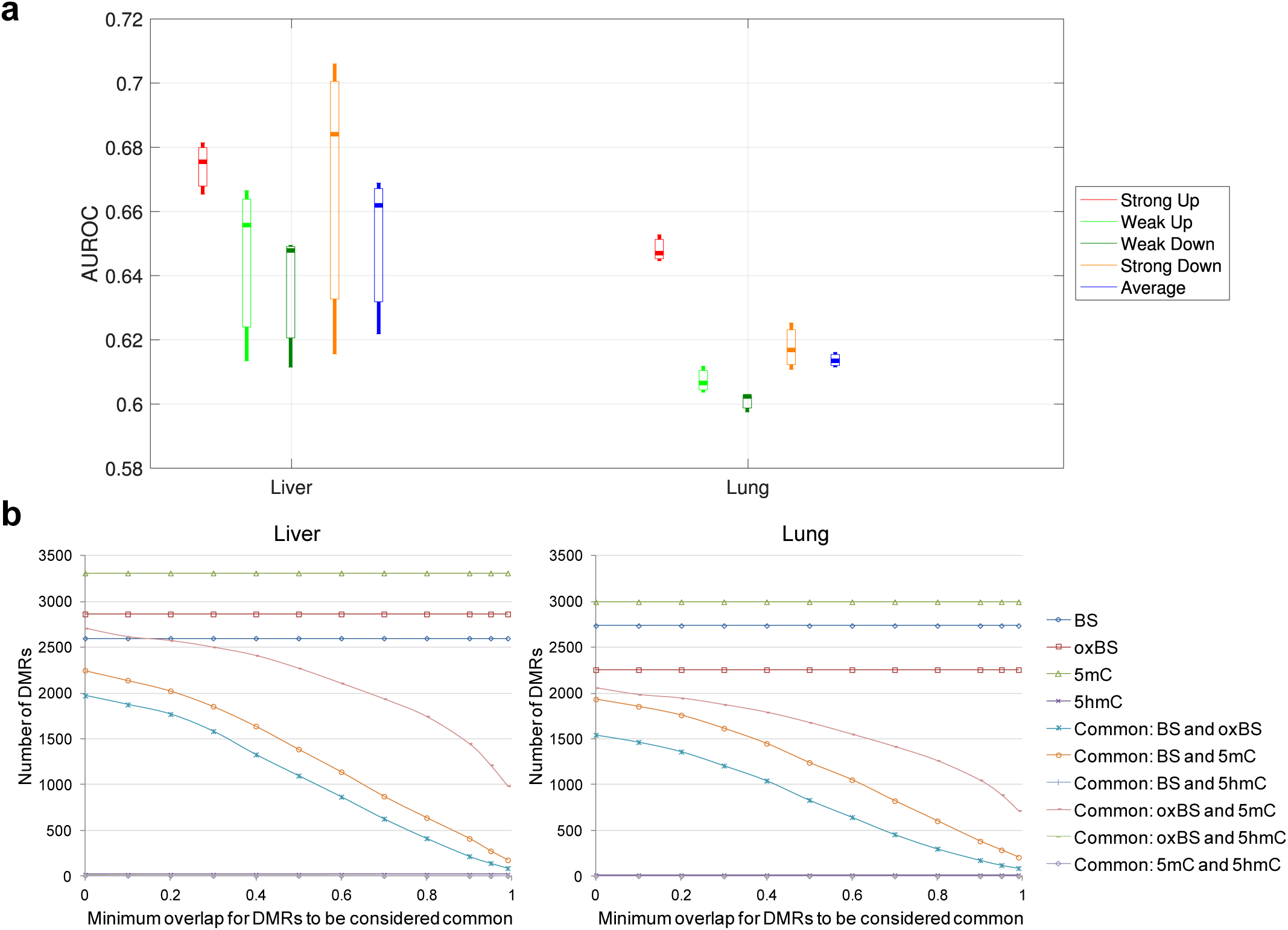
Relationship between methylation and differential expression in cancer. (a) Accuracy of the models for inferring differential expression class, involving all beta value features, based on the large data set with an inter-class gap percentage of 80%. Each bar represents the AUROC values of the three pairs of samples in the group. (b) Overlap of DMRs identified using only WGBS data, only oxWGBS data, only 5mC levels, or only 5hmC levels, for liver (left) and lung (right) samples.

In the above models, the methylation features in the individual samples were used to infer the differential expression class. Another common way to analyze cancer methylome and transcriptome data is to determine DMRs among the tumor and normal samples and look for differentially expressed transcripts potentially caused by them. To evaluate how this standard analysis procedure might be affected by the mixture of 5mC and 5hmC levels in the data, we determined DMRs genome-wide using only BS data, only oxBS data, only 5mC levels, or only 5hmC levels (Materials and Methods). When comparing the overlap of these four sets of DMRs at different stringency thresholds, we found them to differ substantially (Figure 4b). First, we noticed that almost no DMRs were identified based on 5hmC levels, indicating that the 5hmC levels were not sufficiently different between the tumor and normal groups to be considered statistically significant DMRs by the standard DMR calling method. Considering the other three types of data, 5mC consistently gave the highest number of DMRs in both liver and lung samples, suggesting that as compared to the mixture of 5mC and 5hmC signals in BS data, the inferred “clean” 5mC levels were more capable of capturing differential methylation events. Comparing the DMRs identified from BS, oxBS and 5mC, if two DMRs were considered the same as long as their genomic locations had a little bit overlap (minimum overlap ratio close to 0), more than 90% of the oxBS DMRs were also identified from the 5mC data, while DMRs identified from standard BS data could only cover 60-75% of the DMRs identified from oxBS data or 5mC levels. On the other hand, when two DMRs were considered the same only if they had substantial overlaps (with a large overlap ratio), few DMRs identified from these different types of data remained in common. These results show that standard DMR analysis is strongly affected by the type of methylation data involved.

## Discussion

In this study, we have found that 5mC and 5hmC signals provide non-redundant information about gene expression. The median Pearson correlation between the actual log expression levels and the expression levels inferred from methylation data was increased by 18.2% by having separate 5mC and 5hmC signals as compared to having only standard bisulfite sequencing data. Whether this amount of extra information about expression variability is worth the extra cost of producing the additional experimental data (such as oxidative bisulfite sequencing) is a practical decision to be made when designing methylome studies. On the other hand, if the study goal is to identify strongly differentially methylated regions between tumor and normal samples and associate them with differential expression events, our results suggest that it is necessary to have separate measurements of 5mC and 5hmC signals since the DMRs identified from WGBS data could only cover 60-75% of the DMRs identified from “pure” 5mC signals.

In the investigation of PDR features, we found that they were not more informative than beta values in indicating transcript expression levels based on the tumor and normal samples we studied. One limitation of this comparison is that PDR values could only be computed reliably when there were a reasonable number of CpG sites appearing on the same sequencing read and the whole region was covered by a reasonable number of reads. These requirements made the number of transcripts qualified for inclusion very small, since for many transcripts PDR values could not be computed in at least some of the 16 associated genomic regions. An additional difficulty of studying PDR values is that they can only be computed from the raw reads but not from the derived 5mC and 5hmC levels, since the correlation between different CpG sites on the same reads would be loss during the process. It would be useful to further check the usefulness of PDR features in inferring expression levels using additional data sets.

Results in the current study also confirm our previous finding (Lou et al., 2014) that transcript body methylation features are more indicative of gene expression level than promoter methylation features. The highly consistent results from the two studies is remarkable because they involved very different analysis details, including the way of quantifying methylation levels, the cross-validation procedures, the use of gene or transcript as the basic unit, and whether regression of log expression levels is performed. Based on these results, we strongly recommend that when DNA methylation data are used to study transcriptional regulation, methylation signals in the gene body should be included in the analysis, especially the signals at the first exon, last exon and internal introns.

In this study, we investigated methylation of each transcript based on its promoter, body and immediate downstream regions. It would be interesting to extend the study to include enhancer and other distal regulatory elements. Recently, a number of methods have been proposed for identifying target genes of enhancers in a cell type-specific manner (Cao et al., 2017; Corradin et al., 2014; He et al., 2014; Roy et al., 2015; Whalen et al., 2016). However, the accuracy of these methods for cell types without genome contact data such as Hi-C or ChIA-PET is still not high enough for constructing models that can infer expression reliably. The exploration of the general quantitative relationships between enhancer methylation and gene expression levels will need to wait for the availability of more genome contact data or more accurate enhancer-target identification methods.

It has been shown that 5hmC levels are highly variable among tissue types (Nestor et al., 2012). Limited by data availability, we have only included liver and lung samples in our study. Whether 5hmC levels are more indicative of transcript expression levels in other tissue types and whether more DMRs can be identified based on 5hmC levels in other cancer types are questions to be answered when more genome-wide 5mC and 5hmC measurements become available.

## Materials and Methods

### Construction of the data sets

We downloaded raw sequencing read files (.sra) and alignment files (.bam) of the WGBS, oxWGBS and RNA-seq data from Short Read Archive (GSE70091, sub-series GSE70089 for RNA-seq alignment files and GSE70090 for WGBS and oxWGBS raw read files) using the SRA toolkit (https://www.ncbi.nlm.nih.gov/sra/docs/sradownload/). The original data set contained four normal-tumor pairs of liver, but since only three of them had the corresponding RNA-seq data, we excluded this fourth pair from all our analyses. Following Li et al. (2016), we aligned the WGBS and oxWGBS raw reads to the human reference genome hg19 using BSmooth (Hansen et al., 2012). Read pairs having identical alignments of both mates were considered potential duplicates due to PCR artifacts, and only one read pair was retained for each set of duplicate read pairs.

For each CpG site, we computed the beta value of it as the number of reads supporting an unconverted cytosine divided by the total number of reads covering the site, for both WGBS and oxWGBS data sets. To ensure the reliability of the input data, following the data processing in Li et al. (2016), we excluded reads with a mapping quality less than 20, bases on a read with a base quality less than 10, and bases within the 10 5’-most positions of both mates of each read pair. To reduce effects of sampling errors, CpG sites with fewer than 5 aligned reads were also excluded.

We further computed 5mC and 5hmC levels of each site using a maximum likelihood method (Xu et al., 2016). For each of the four types of methylation features, these per-site methylation levels were then aggregated to compute the average level of each associated region defined for a transcript. These associated regions included 16 regions overlapping or immediately next to the transcript (Lou et al., 2014), namely five consecutive 400bp bins upstream of the transcription start site (TSS) (Up1-Up5, with Up1 closest to the TSS), first exon (FirstEx), first intron (FirstIn), internal exons (IntEx), internal introns (IntIn), last exon (LastEx), last intron (LastIn), and five consecutive 400bp bins downstream of the transcription termination site (TTS) (Down1-Down5, with Down1 closest to the TTS). If any associated region had fewer than 5 CpG sites with at least 5 aligned reads, the whole transcript was discarded. By default, we included all annotated protein-coding and non-coding transcripts of levels 1, 2 and 3 in Gencode (Harrow et al., 2012) version 19, while in some analyses we only considered a subset of these transcripts to see how the results differed.

In order to have all 16 regions defined, we always considered only transcripts with at least four exons and only transcripts. We also discarded transcripts having less than 3 CpG sites with at least 5 aligned reads any of the 16 regions.

In addition, for each region associated to a gene, we calculated its PDR value in each sample as the ratio of reads having discordant methylation status. Specifically, we considered reads with an alignment that overlapped the region, with reads having a mapping quality less than 20 or covering less than 3 CpG sites excluded. For each of the remaining reads, it was considered a concordant read if less than 10% or more than 90% of the CpG sites it covered had the same methylation status. The other reads were considered discordant, and the number of them was used to compute the PDR value. To ensure the robustness of the computed PDR values, transcripts with any one of the 16 associated regions having less than 3 aligned reads were discarded. This filtering was the main reason that the resulting small data set had a much smaller number of transcripts than the big data set, which only had beta values as features. Since the calculations of PDR values required information of sequencing reads rather than individual CpG sites, they could not be computed for the processed 5mC and 5hmC data sets, which did not have read-level information anymore.

We also computed transcript expression levels, defined as fragments per kilobase per million mapped reads (FPKM), using Cufflinks (Trapnell et al., 2010) version 2.2.1 using the -G option.

### Statistical modeling of expression classes

In each sample, we defined a high-expression class of transcripts as those having an FPKM value of 1 or more, a low-expression class of transcripts as those having an FPKM value larger than 1E-10 but smaller than 1E-2, and a zero-expression class of transcripts as those having an FPKM value less than 1E-100.

We modeled the expression class using either all or a subset of the methylation features. We chose Random Forest models (Bagging with 50 random trees as the base classifiers) since they were previously shown to perform well for modeling the quantitative roles of DNA methylation (Lou et al., 2014).

We designed a cross-validation procedure for evaluating the performance of the models as follows. We paired up short and long autosomes, namely chromosome 1 with chromosome 22, chromosome 2 with chromosome 21, and so on, leading to 11 chromosome pairs. Each time one of the chromosome pairs was left out for testing, while the other 10 pairs were used for training a model. The model was then applied to the transcripts in the left-out chromosome pair, either from the same sample (within-sample test) or from another sample (across-sample test). Finally, the predictions from the 11 left-out sets were combined to compute the performance metric AUROC (area under the receiver-operator characteristics). This design of the cross-validation procedure avoids two types of trivial memorization. First, in the within-sample test, if all transcripts were randomly distributed to the training and testing sets instead, two transcripts from the same gene could be respectively assigned to the training and testing sets, leading to a simple memorization of the training transcript’s expression level when predicting the testing transcript’s expression level, since they would share very similar methylation features. Second, in the between-sample test, if all transcripts from a sample was used to construct the model instead, when applying the model to another sample, again the predictions could be simply memorization of the expression levels of the same transcripts in the training sample, when the training and testing samples were highly similar, such as those from the same tissue type and disease state. In addition to avoiding such memorization, the pairing of long and short chromosomes in our procedure also led to a similar number of transcripts in each chromosome pair. To make our results more reliable, we also repeated each classification task 10 times with different random seeds used to construct the Random Forest models, and reported the average performance.

### Feature importance evaluation

To evaluate the importance of different features in explaining transcript expression levels, we used a forward-searching procedure to construct models with only subsets of the most useful features. Specifically, we started with constructing models having only one feature, and compared their performance. The feature used in the most accurate model was then added to the set of selected features, and new models were constructed by having this feature plus one of the remaining features. This procedure was repeated iteratively, with the feature leading to the best performance in each iteration added to the set of selected features. The whole procedure ended when all features had been selected. Finally, we gave the first *x* features selected a score of *x, x*-1,…, 1, respectively, where *x* was chosen to be 1/2 of the total number of features when all 64 features were considered separately, and 2/3 of the total number of features/feature blocks in all other settings.

We performed this feature importance evaluation with each sample, and also summed the scores across all samples to define a single importance score for each feature.

In addition to individual features, we also used this procedure to evaluate the importance of different feature blocks.

### Statistical modeling of expression levels

Since the grouping of transcripts into expression classes relied on a specified way of defining the classes and a specific number of classes, to ensure that our findings were not affected by our specific choices, we also constructed regression models to predict the log expression level of the transcripts based on their methylation features. Specifically, for a transcript with a FPKM value of *y*, we used log_10_(*y* + 1) as the prediction target. We used support vector regression with a radial basis function (RBF) kernel to construct the models. Model performance was evaluated using the same cross-validation procedure as in the case of predicting expression classes, quantified by both Pearson correlation and Spearman correlation values.

### Definition of differential expression classes

To define differential expression classes, we first calculated a differential expression level of each transcript by subtracting its FPKM value in a normal sample from its FPKM value in the corresponding tumor sample, FPKMdiff = FPKMtumor - FPKMnormal. We kept only transcripts with no missing data in all 6 samples pairs. Then for each tissue type, we used the median differential expression level of a transcript among the three sample pairs to determine its differential expression class. Transcripts with a median differential expression level between —10^−5^ and 10^−5^ were considered having no significant change of expression and were not included in any of the classes. For the remaining transcripts with a positive differential expression level (i.e., higher expression in tumor), we took the top and bottom *x%* of transcripts with the largest and smallest absolute differential expression values to define the strongly up-regulated and weakly up-regulated classes, respectively, where *x* is a variable and we called 1 — 2*x%* as the gap percentage between the two classes. We tried different values of the gap percentage from 10 to 90, with 80 used as the default as a tradeoff between the clear separation of the two classes and the number of transcripts that can be included in them. In the same way, we also defined the strongly down-regulated and weakly down-regulated classes.

### Statistical modeling of differential expression classes

We compared the sizes of the four classes, and randomly down-sampled the larger ones until all four classes had the same number of transcripts. This random down-sampling was repeated 10 times to generate 10 different data sets. We then trained and tested Random Forest models for the 4 differential expression classes in a one-against-all manner using the same cross-validation procedure as in the case of modeling expression classes.

### Analysis of differentially methylated regions

Differentially methylated regions (DMRs) were identified by Metilene v0.2-6 (Jühling et al., 2016) based on the beta values of CpG sites from the four types of methylation data. For each tissue type, the three tumor samples were compared with the three normal samples to identify the DMRs. We further filtered the DMRs by retaining only those at least 100bp long with at least 3 CpG sites having 10x read coverage, and an average difference of beta value between the samples in the tumor and non-tumor groups at least 0.1. We also tried 5 other sets of values for these filtering parameters, but the resulting trends were all highly similar. We considered a DMR called from one data set to overlap a DMR called from another data set if the intersection of them constitutes at least *x*% of both DMRs, where *x* is the minimum overlap ratio and we tried a range of values for it. To count the number of DMRs commonly called from two data sets, we first obtained two numbers, namely the number of DMRs in the first set that overlaps one or more DMRs in the second set, and the number of DMRs in the second set that overlaps one or more DMRs in the first set. It turns out that these two sets of numbers were usually identical and differed at most by a small number. We therefore used their average in our report.

## Acknowledgments

QW, ASLC and KYY were partially supported by HKSAR RGC CRF C4017-14G.

